# Failure to detect synergy between variants in transferrin and hemochromatosis and Alzheimer’s disease in large cohort

**DOI:** 10.1101/649962

**Authors:** Elizabeth Vance, Josue Gonzalez, Justin B. Miller, Alzheimer’s Disease Genetics Consortium (ADGC), Lindsey Staley, Paul K. Crane, Shubhabrata Mukherjee, John S.K. Kauwe

## Abstract

Alzheimer’s disease (AD) is the most common cause of dementia and, despite decades of effort, there is no effective treatment. In the last decade, many association studies have identified genetic markers that are associated with AD status. Two of these studies suggest that an epistatic interaction between variants *rs1049296* in the Transferrin (*TF*) gene and *rs1800562* in the Homeostatic Iron Regulator (*HFE)* gene, commonly known as “the hemochromatosis gene”, is in genetic association with AD. *TF* and *HFE* are involved in the transport and regulation of iron in the brain, and disrupting these processes exacerbates AD pathology through increased neurodegeneration and oxidative stress. However, by using a significantly larger dataset from the Alzheimer’s Disease Genetics Consortium (ADGC), we fail to detect an association between *TF rs1049296* or *HFE rs1800562* with AD risk (*TF rs1049296* p=0.38 and *HFE rs1800562* p=0.40). In addition, logistic regression with an interaction term and a Synergy Factor Analysis (SFA) both failed to detect epistasis between *TF rs1049296* and *HFE rs1800562* (SF=0.94; p=0.48) in AD cases. Each of these analyses had sufficient statistical power (Power>0.99), suggesting that previously-reported associations may be the result of more complex epistatic interactions, genetic heterogeneity, or were false-positive associations due to limited sample sizes.

## Introduction

Alzheimer’s disease (AD) is the most common cause of dementia and inflicts an estimated 24 to 35 million people worldwide, with incidences predicted to increase dramatically as the population ages (“2018 Alzheimer’s disease facts and figures,” 2018). Although decades of research have been spent investigating the causes and architecture of this neurodegenerative disease, it still inflicts an estimated 5.7 million people in the United States alone. This number is projected to increase to 13.8 million by mid-century (“2018 Alzheimer’s disease facts and figures,” 2018). Association studies have accurately identified single-nucleotide polymorphisms (SNPs) associated with AD (D. Harold et al., 2009; Denise Harold et al., 2009; Hollingworth et al., 2011; J.-C. Lambert et al., 2009; J. C. Lambert et al., 2013; Seshadri et al., 2010; Shen et al., 2015; Shuai et al., 2015; Yan et al., 2015). However, these genetic loci account for only a fraction of AD heritability, (Ridge, Mukherjee, Crane, Kauwe, & Alzheimer’s Disease Genetics, 2013) suggesting that much of AD’s unexplained genetic make-up may be due to epistasis (Bullock et al., 2013; Combarros, Cortina-Borja, Smith, & Lehmann, 2009; M. T. Ebbert et al., 2014; Infante et al., 2004). Epistasis occurs when multiple genes interact to create a single phenotype (Cordell, 2002). These kinds of synergetic relationships play a critical role in the etiology of complex diseases, yet remain vastly understudied in AD pathology (“2018 Alzheimer’s disease facts and figures,” 2018; M. T. W. Ebbert, Ridge, & Kauwe, 2015; Raghavan & Tosto, 2017).

The Transferrin (*TF*) gene and the Homeostatic Iron Regulator (*HFE)* gene, commonly known as “the hemochromatosis gene”, have been reported to show epistasis and play a role in the development of AD (Robson et al., 2004; Tisato et al., 2018). TFs are a group of non-heme iron-binding glycoproteins found in fluids and cells of vertebrates. The main role of *TF* is to maintain iron homeostasis in the body (Gkouvatsos, Papanikolaou, & Pantopoulos, 2012). In the brain, *TF* interacts with the Amyloid Precursor Protein (APP) (Belaidi et al., 2018) and tau (Jahshan, Esteves-Villanueva, & Martic-Milne, 2016), two of the major protein families implicated in AD pathology. Since iron is essential for oxygen transport, its mis-regulation in the brain can lead to oxidative stress and neurodegeneration (Dias, Junn, & Mouradian, 2013; Matak et al., 2016; Yarjanli, Ghaedi, Esmaeili, Rahgozar, & Zarrabi, 2017). *HFE* encodes for a transmembrane glycoprotein that binds to a *TF* receptor, subsequently regulating iron in the cell (Bennett, Lebron, & Bjorkman, 2000; Feder et al., 1996; Lebron et al., 1998). Mutations in *HFE* are associated with neurodegenerative diseases through increasing neuroinflammation and production of free radicals in the brain (Andersen, Johnsen, & Moos, 2014; Lull & Block, 2010). In addition, other studies suggest that *TF* and *HFE* are involved in the transport and regulation of iron in the brain, and disrupting these processes potentially affects AD pathology through increased neurodegeneration and oxidative stress (Ali-Rahmani, Schengrund, & Connor, 2014; Lehmann et al., 2006).

Robson et al. (2004) suggested that epistasis between *TF* variant *rs1049296* and *HFE* variant *rs1800562* is associated with AD. Although neither SNP alone was a risk factor for AD, the presence of both alleles resulted in a five times greater risk of developing AD. (Robson et al., 2004). Since the sample size for that study was relatively small (191 cases and 269 controls), a replication of these findings on a slightly larger dataset (1,161 cases and 1,342 controls) was conducted and corroborated a significant association with AD risk among bi-allelic carriers of *rs1049296* and *rs1800562* (Kauwe et al., 2010).

Our study expands on these previous studies and attempts to detect statistical epistasis between *TF rs1049296* and *HFE rs1800562* with respect to AD risk using 25,666 individuals from the Alzheimer’s Disease Genetics Consortium (ADGC), which is an expansion of the dataset employed by Kauwe et al. (2010).

## Material and Methods

### Dataset and Filtering

Our analysis started with GWAS data from all 28,730 individuals in the Alzheimer’s Disease Genetic Consortium (ADGC) dataset as described by Naj et al. (A. C. Naj et al., 2011). ADGC is a collection of 30 merged datasets spanning 1984 to 2012, and was established to help identify genetic markers of late onset AD. (Boehme, Mukherjee, Crane, & Kauwe, September 2014). ADGC imputed the 30 datasets to the Haplotype Reference Consortium (HRC) reference panel, which includes 64,976 haplotypes and 39,235,157 SNPs (Loh et al., 2016; Adam C. Naj et al., 2017). Genotyped markers with a minor allele frequency less than 1% and a deviation from Hardy Weinberg Equilibrium (HWE) where α<10^-6^ were removed. All aspects of the study were approved by institutional review boards, and each applicant signed a written form of consent for their genetic data to be used for research purposes.

We followed the same filtering protocols established by Ridge et al. (Ridge et al., 2013) by genotyping markers with a minor allele frequency less than 1% and removing markers with a HWE p-value less than 10^-6^. Principle components were calculated using Eigensoft (Patterson, Price, & Reich, 2006; Price et al., 2006) to account for population specific variations in allele distribution. After filtering, 12,532 cases and 13,134 control subjects contained genotypic information for *TF rs1049296* and *HFE rs1800562*.

### Genetic Analyses

The main effects of *TF rs1049296* and *HFE rs1800562* on AD risk were measured using a multivariate nonparametric logistic regression analysis. Each SNP was first analyzed as a single term and then as an interaction term in a subsequent analysis. We included sex, age of onset, *APOE e4* allele status, AD status, and 10 principle components as covariates in our analysis. In addition, we performed a chi-square analysis to determine odds ratios between AD status in each SNP as a single term and as an interaction term, respectively. Lastly, we performed a Synergy Factor Analysis (SFA) (Cortina-Borja, Smith, Combarros, & Lehmann, 2009). These analyses were performed for each of the 30 cohorts separately and for the entire ADGC dataset combined as a single cohort.

Furthermore, we calculated the power of analysis for the ADGC dataset using G*Power (Faul, Erdfelder, Lang, & Buchner, 2007). The computations for power of the previous analysis performed by Kauwe et al. (2010) revealed that for a sample size of 2,503 and an alpha of 0.05, their logistic regression model had power of 0.95 to detect an effect size of 0.86. The power of our analysis reveals that for a sample size of 25,666 and the same alpha of 0.05, our logistic regression model has power of >0.99 to detect a similar effect size of 0.86.

## Results

The nonparametric logistic regression analysis using ADGC as one cohort demonstrated that when testing the main effects, neither *TF rs1049296* nor *HFE rs1800562* was associated with AD risk (*TF rs1049296* p=0.38; *HFE rs1800562* p=0.40). The logistic regression analyses including an interaction term for the two variants also failed to show significant association (p=0.23). Similarly, the SFA analysis did not find epistasis between *TF rs1049296* and *HFE rs1800562* (SF=0.94; p=0.48).

We performed logistic regression on all 30 individual cohorts (see Figure 1). We detected a significant epistatic association between the interaction term and AD status in the ACT cohort (p=0.038) and a suggested association in the ADC1 cohort (p=0.063). In addition, the individual effect of *HFE rs1800562* shows a suggested association with AD status in the ADC6 (p=0.099), WHICAP (p=0.052), ADC4 (p=0.076), and ROSMAP (p=0.094) cohorts. Furthermore, logistic regression for the individual effect of *TF rs1049296* determined a significant association with AD status in the WASHU cohort (p=0.016). However, none of these associations remained significant after a Bonferroni correction for multiple tests.

**Figure 1:**
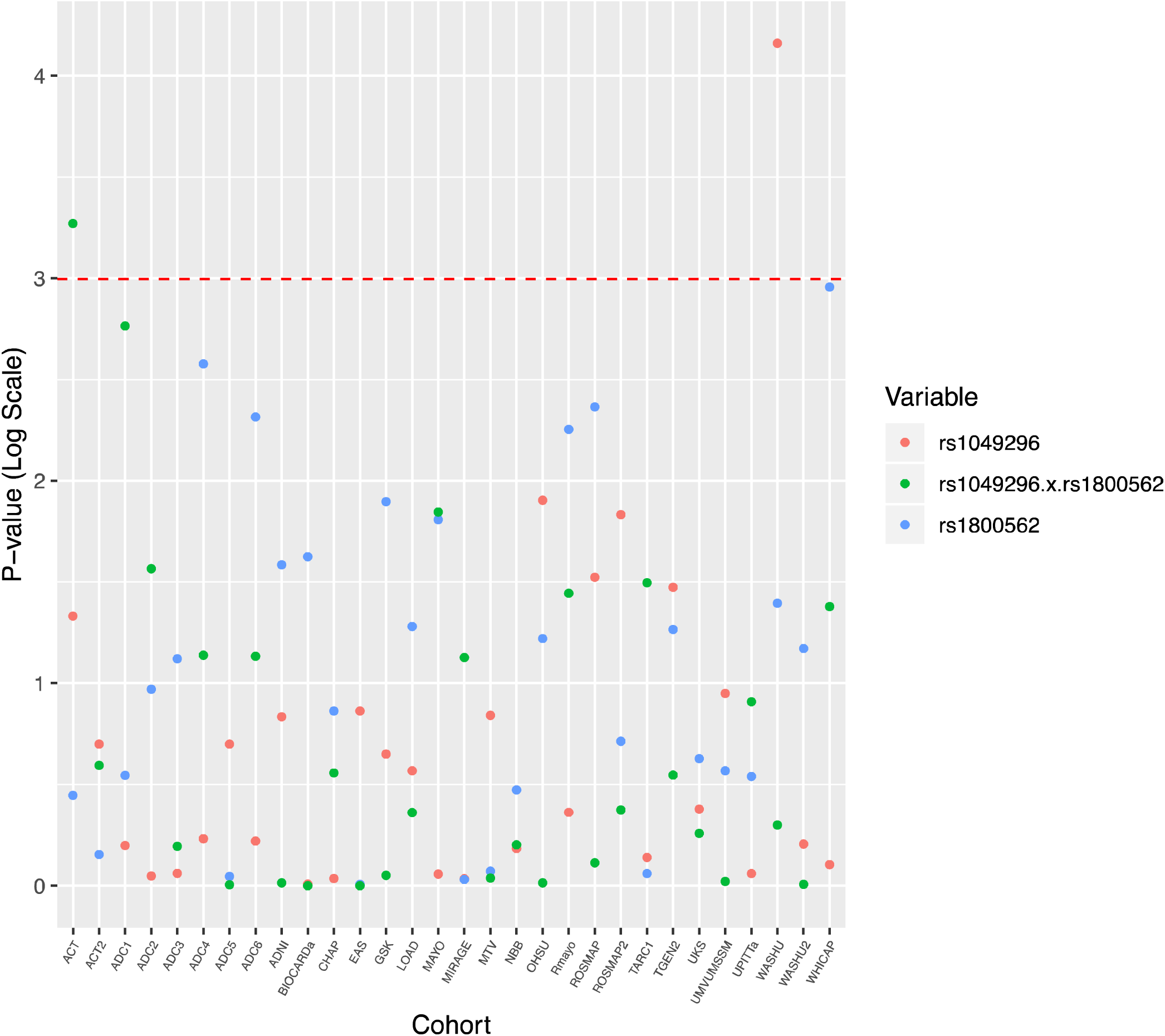
Logistic Regression P-values per Cohort. We performed logistic regression on each cohort to determine the p-values for rs1049296, rs1800562, and the epistatic interaction of these variants. Each cohort is shown on the x-axis, and the p-value for each cohort is shown on the y-axis.The red line indicates the alpha value of 0.05. From our analysis, only the ACT and WASHU cohorts have significant p-values at these variants.

In addition, chi-squared analyses between terms and AD status demonstrated a non-significant likelihood for any single term or interaction. The odds ratio for *rs1049269* was 0.97 with a 95% confidence interval (CI) between 0.92 and 1.03, while *rs1800562* had an odds ratio of 1.06 with a CI of 0.98 to 1.15, and the interaction term had an odds ratio of 0.99 with a CI of 0.86 to 1.14. The odds ratios and confidence intervals for main effects and the interaction in each cohort are displayed in Figure 2.

**Figure 2:**
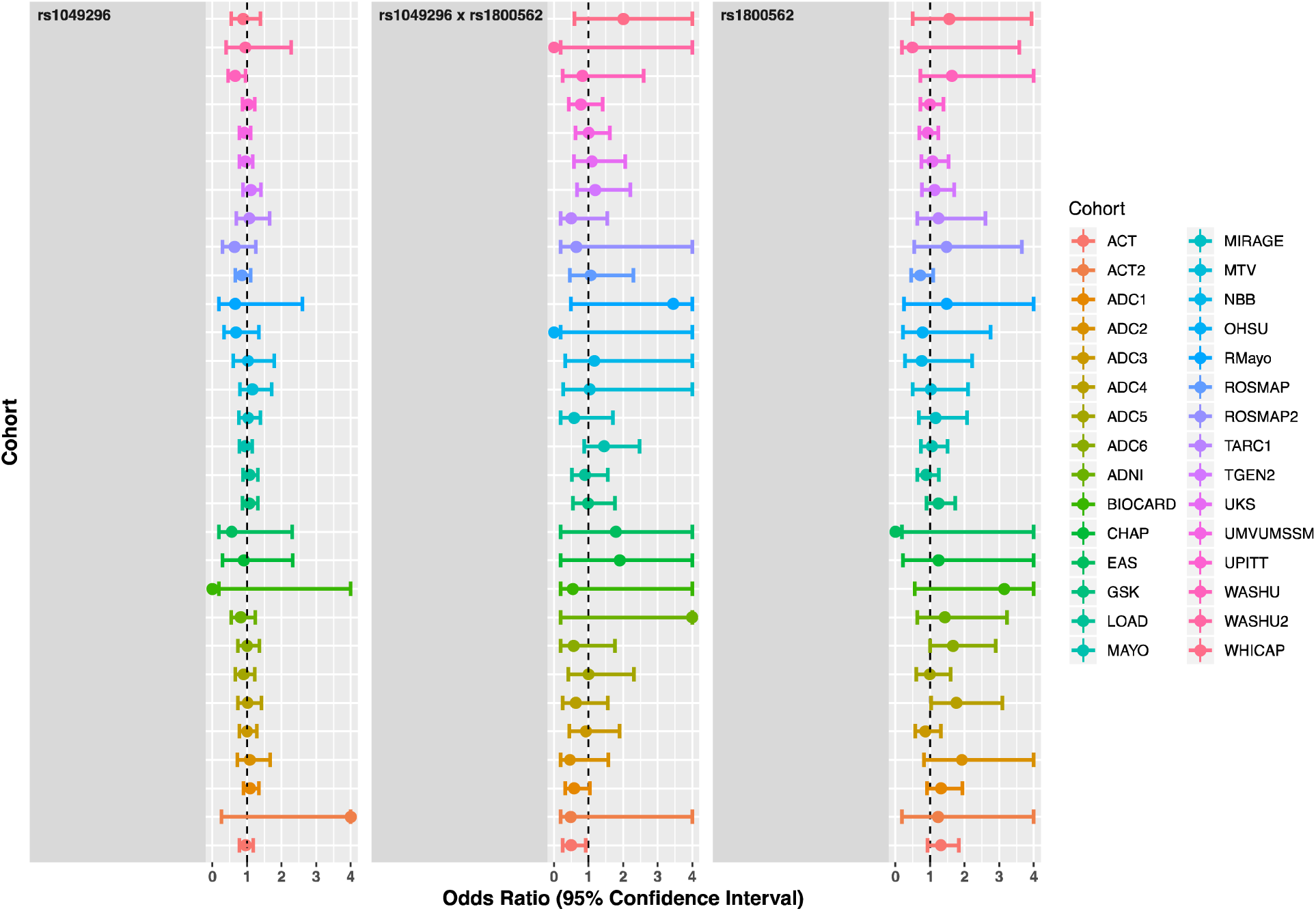
Logistic Regression odds Ratios and Confidence Intervals per Cohort. We performed logistic regression on each cohort to determine the odds ratios for rs1049296,rs1800562, and the epistatic interaction of these variants. Each cohort is shown on the y-axis. and the odds radio with the 95% confidence interval for each cohort is shown on the x-axis. Lower limit values are truncated to .2, while upper limit and odds ratio values are truncated to 4. The dashed line indicates an odds ratio value of 1. Although the ACT2,ADNI,BIOCARD,Rmayo, and WHICAP cohorts have seemingly high odds ratio values (>2), it is important to note that the confidence intervals for these cohorts span the null value (1) and are not precise. Furthermore the p-values for these cohorts suggest that the effects are not significant (see Figure 1)

## Discussion

We failed to detect evidence of epistasis between *TF rs1049296* and *HFE rs1800562* as a risk for AD in the ADGC dataset. These findings do not support the conclusions drawn in the previous reports by Robson et al. (2004) and Kauwe et al. (Kauwe et al., 2010). The cause for this variability among studies could be a result of genetic heterogeneity, the complex nature of epistasis, or false positives in these previous studies due to limited sample size.

Although recent literature suggests that much of the unidentified genetic makeup of AD is due to epistasis (Bullock et al., 2013; Combarros et al., 2009; M. T. Ebbert et al., 2014; Infante et al., 2004; Mez, 2016), the complex nature of these gene-gene interactions makes it difficult for them to be accurately measured and defined (Gilbert-Diamond & Moore, 2011; Kouyos, Silander, & Bonhoeffer, 2007; Urbanowicz, Kiralis, Fisher, & Moore, 2012). Models for epistatic interactions are challenging to create because the models require large datasets to analyze combinations of variables simultaneously (Moore & Williams, 2009). When an insufficient number of samples are used, results have poor statistical power, which leads to frequent false negatives in gene-gene interaction studies. Likewise, the numerous comparisons required to assess epistasis may generate false positive findings (Mackay & Moore, 2014). Inadequate sample size can also result in false positives and is identified through statistical power analyses (Christley, 2010). The experiments performed by Robson et al. (2004) and Kauwe et al. (Kauwe et al., 2010) used datasets with much fewer individuals than the dataset used in this manuscript, and consequently have lower statistical power than our analysis. Although Kauwe et al. (2010) appeared to have sufficient power for their study (0.95), it is still possible that their results were false positive findings. Current research suggests a troubling phenomenon known as the “winner’s curse,” which occurs when the estimated effect of an association is inflated compared to the true genetic effect and the effects later measured in follow-up studies (Huang, Ritchie, Brozynska, & Inouye, 2018; Palmer & Pe’er, 2017). The level of power necessary to accurately detect epistasis is currently unknown, and as such, replication studies are a necessary part of validating epistasis. As our results show, even studies that appear to have sufficient power, such as Kauwe et al. (2010), should be re-evaluated when larger datasets become available.

Heterogeneity in the genetic causes of AD is certainly present (Mez, 2016), and further erodes power to detect statistical epistasis. Finally, even when statistical evidence for epistasis is detected, it does not necessarily indicate the presence of a physical biological interaction between the implicated proteins (M. T. W. Ebbert et al., 2015). Statistical patterns can be a product of a variety of underlying mechanisms. Therefore, the complexity of biological and statistical epistasis could also account for disparities in replication studies. Increasing sample sizes gives us better statistical power. Likewise, increasing the amount of multidimensional - omics data will help us focus our efforts on specific candidate interactions. We anticipate that as more multidimensional -omics data become available, our ability to understand the role of epistasis in AD risk will improve and help in the development of novel approaches to prevent and treat the disease.

## Acknowledgements

The National Institutes of Health, National Institute on Aging (NIH-NIA) supported this work through the following grants: ADGC, U01 AG032984, RC2 AG036528; Samples from the National Cell Repository for Alzheimer’s Disease (NCRAD), which receives government support under a cooperative agreement grant (U24 AG21886) awarded by the National Institute on Aging (NIA), were used in this study. We thank contributors who collected samples used in this study, as well as patients and their families, whose help and participation made this work possible; Data for this study were prepared, archived, and distributed by the National Institute on Aging Alzheimer’s Disease Data Storage Site (NIAGADS) at the University of Pennsylvania (U24-AG041689-01); NACC, U01 AG016976; NIA LOAD (Columbia University), U24 AG026395, U24 AG026390, R01AG041797; Banner Sun Health Research Institute P30 AG019610; Boston University, P30 AG013846, U01 AG10483, R01 CA129769, R01 MH080295, R01 AG017173, R01 AG025259, R01 AG048927, R01AG33193, R01 AG009029; Columbia University, P50 AG008702, R37 AG015473, R01 AG037212, R01 AG028786; Duke University, P30 AG028377, AG05128; Emory University, AG025688; Group Health Research Institute, UO1 AG006781, UO1 HG004610, UO1 HG006375, U01 HG008657; Indiana University, P30 AG10133, R01 AG009956, RC2 AG036650; Johns Hopkins University, P50 AG005146, R01 AG020688; Massachusetts General Hospital, P50 AG005134; Mayo Clinic, P50 AG016574, R01 AG032990, KL2 RR024151; Mount Sinai School of Medicine, P50 AG005138, P01 AG002219; New York University, P30 AG08051, UL1 RR029893, 5R01AG012101, 5R01AG022374, 5R01AG013616, 1RC2AG036502, 1R01AG035137; North Carolina A&T University, P20 MD000546, R01 AG28786-01A1; Northwestern University, P30 AG013854; Oregon Health & Science University, P30 AG008017, R01 AG026916; Rush University, P30 AG010161, R01 AG019085, R01 AG15819, R01 AG17917, R01 AG030146, R01 AG01101, RC2 AG036650, R01 AG22018; TGen, R01 NS059873; University of Alabama at Birmingham, P50 AG016582; University of Arizona, R01 AG031581; University of California, Davis, P30 AG010129; University of California, Irvine, P50 AG016573; University of California, Los Angeles, P50 AG016570; University of California, San Diego, P50 AG005131; University of California, San Francisco, P50 AG023501, P01 AG019724; University of Kentucky, P30 AG028383, AG05144; University of Michigan, P50 AG008671; University of Pennsylvania, P30 AG010124; University of Pittsburgh, P50 AG005133, AG030653, AG041718, AG07562, AG02365; University of Southern California, P50 AG005142; University of Texas Southwestern, P30 AG012300; University of Miami, R01 AG027944, AG010491, AG027944, AG021547, AG019757; University of Washington, P50 AG005136, R01 AG042437; University of Wisconsin, P50 AG033514; Vanderbilt University, R01 AG019085; and Washington University, P50 AG005681, P01 AG03991, P01 AG026276. The Kathleen Price Bryan Brain Bank at Duke University Medical Center is funded by NINDS grant # NS39764, NIMH MH60451 and by Glaxo Smith Kline. Support was also from the Alzheimer’s Association (LAF, IIRG-08-89720; MP-V, IIRG-05-14147), the US Department of Veterans Affairs Administration, Office of Research and Development, Biomedical Laboratory Research Program, and BrightFocus Foundation (MP-V, A2111048). P.S.G.-H. is supported by Wellcome Trust, Howard Hughes Medical Institute, and the Canadian Institute of Health Research. Genotyping of the TGEN2 cohort was supported by Kronos Science. The TGen series was also funded by NIA grant AG041232 to AJM and MJH, The Banner Alzheimer’s Foundation, The Johnnie B. Byrd Sr. Alzheimer’s Institute, the Medical Research Council, and the state of Arizona and also includes samples from the following sites: Newcastle Brain Tissue Resource (funding via the Medical Research Council, local NHS trusts and Newcastle University), MRC London Brain Bank for Neurodegenerative Diseases (funding via the Medical Research Council),South West Dementia Brain Bank (funding via numerous sources including the Higher Education Funding Council for England (HEFCE), Alzheimer’s Research Trust (ART), BRACE as well as North Bristol NHS Trust Research and Innovation Department and DeNDRoN), The Netherlands Brain Bank (funding via numerous sources including Stichting MS Research, Brain Net Europe, Hersenstichting Nederland Breinbrekend Werk, International Parkinson Fonds, Internationale Stiching Alzheimer Onderzoek), Institut de Neuropatologia, Servei Anatomia Patologica, Universitat de Barcelona. ADNI data collection and sharing was funded by the National Institutes of Health Grant U01 AG024904 and Department of Defense award number W81XWH-12-2-0012. ADNI is funded by the National Institute on Aging, the National Institute of Biomedical Imaging and Bioengineering, and through generous contributions from the following: AbbVie, Alzheimer’s Association; Alzheimer’s Drug Discovery Foundation; Araclon Biotech; BioClinica, Inc.; Biogen; Bristol-Myers Squibb Company; CereSpir, Inc.; Eisai Inc.; Elan Pharmaceuticals, Inc.; Eli Lilly and Company; EuroImmun; F. Hoffmann-La Roche Ltd and its affiliated company Genentech, Inc.; Fujirebio; GE Healthcare; IXICO Ltd.; Janssen Alzheimer Immunotherapy Research & Development, LLC.; Johnson & Johnson Pharmaceutical Research & Development LLC.; Lumosity; Lundbeck; Merck & Co., Inc.; Meso Scale Diagnostics, LLC.; NeuroRx Research; Neurotrack Technologies; Novartis Pharmaceuticals Corporation; Pfizer Inc.; Piramal Imaging; Servier; Takeda Pharmaceutical Company; and Transition Therapeutics. The Canadian Institutes of Health Research is providing funds to support ADNI clinical sites in Canada. Private sector contributions are facilitated by the Foundation for the National Institutes of Health (www.fnih.org). The grantee organization is the Northern California Institute for Research and Education, and the study is coordinated by the Alzheimer’s Disease Cooperative Study at the University of California, San Diego. ADNI data are disseminated by the Laboratory for Neuro Imaging at the University of Southern California. We thank Drs. Dallas Anderson and Marilyn Miller from NIA who are *ex-officio* ADGC members.

